# RefZ and Noc act synthetically to prevent aberrant divisions during *Bacillus subtilis* sporulation

**DOI:** 10.1101/2022.01.13.476237

**Authors:** Allyssa K. Miller, Jennifer K. Herman

## Abstract

During sporulation, *Bacillus subtilis* undergoes an atypical cell division that requires overriding mechanisms which protect chromosomes from damage and ensure inheritance by daughter cells. Instead of assembling between segregated chromosomes at midcell, the FtsZ-ring (Z-ring) coalesces polarly, directing division over one chromosome. The DNA-binding protein RefZ facilitates the timely assembly of polar Z-rings and partially defines the region of chromosome initially captured in the forespore. RefZ binds to motifs (*RBM*s) located proximal to the origin of replication (*oriC*). Although *refZ* and the *RBM*s are conserved across the *Bacillus* genus, a *refZ* deletion mutant sporulates with wildtype efficiency, so the functional significance of RefZ during sporulation remains unclear. To further investigate RefZ function, we performed a candidate-based screen for synthetic sporulation defects by combining Δ*refZ* with deletions of genes previously implicated in FtsZ regulation and/or chromosome capture. Combining Δ*refZ* with deletions of *ezrA, sepF, parA*, or *minD* did not detectably affect sporulation. In contrast, a Δ*refZ* Δ*noc* mutant exhibited a sporulation defect, revealing a genetic interaction between RefZ and Noc. Using reporters of sporulation progression, we determined the Δ*refZ* Δ*noc* mutant exhibited sporulation delays after Spo0A activation but prior to late sporulation, with a subset of cells failing to divide polarly or activate the first forespore-specific sigma factor, SigF. The Δ*refZ* Δ*noc* mutant also exhibited extensive dysregulation of cell division, producing cells with extra, misplaced, or otherwise aberrant septa. Our results reveal a previously unknown epistatic relationship that suggests *refZ* and *noc* contribute synthetically to regulating cell division and supporting spore development.

**IMPORTANCE:** The DNA-binding protein RefZ and its binding sites (*RBM*s) are conserved in sequence and location on the chromosome across the *Bacillus* genus and contribute to the timing of polar FtsZ-ring assembly during sporulation. Only a small number of non-coding and non-regulatory DNA motifs are known to be conserved in chromosomal position in bacteria, suggesting there is strong selective pressure for their maintenance; however a *refZ* deletion mutant sporulates efficiently, providing no clues as to their functional significance. Here we find that in the absence of the nucleoid occlusion factor Noc, deletion of *refZ* results in a sporulation defect characterized by developmental delays and aberrant divisions.

## INTRODUCTION

Chromosome inheritance depends on precise division site selection. Abnormal divisions can result in aneuploidy, including total chromosome loss. Eukaryotes employ cell cycle checkpoints to ensure replication and segregation are complete before cytokinesis initiates. In contrast, bacteria often segregate DNA concurrently with division, so mechanisms to coordinate these processes are critical. To ensure faithful transmission of genetic material to progeny, bacteria segregate replicated DNA to opposite cell halves and divide between chromosome masses (nucleoids). Division at midcell is initiated by polymerization and bundling of membrane-tethered FtsZ protofilaments into the “Z-ring,” a dynamic structure subject to both positive and negative regulation (1-20). The Z-ring facilitates recruitment of additional factors needed for division at midcell, including peptidoglycan remodeling enzymes (2, 14-20).

The Min system and Nucleoid Occlusion (NO) are redundant, but mechanistically distinct systems for ensuring Z-rings assemble at midcell, between chromosomes (21, 22). In *B. subtilis*, the MinCD complex localizes in the immediate vicinity of the nascent septum and inhibits additional Z-rings from forming (23-26); following division, Min inhibition persists at the newly formed, nucleoid-free poles (27, 28). NO, by contrast, prevents assembly of division-competent Z-rings over the bulk of the nucleoid (29, 30). In *E. coli*, NO is mediated by an inhibitor of FtsZ polymerization, SlmA. SlmA is also a DNA-binding protein, with specificity for motifs (SBSs) enriched throughout the chromosome except in the terminus (*ter*) region (30, 31). Following chromosome replication and segregation, *ter* is localized at midcell. The coincident segregation of SlmA away from midcell leads to release of NO, a condition more favorable to, but not sufficient for, Z-ring assembly. The NO protein of *B. subtilis*, Noc, also binds to motifs enriched distal to *ter* (32); however, unlike SlmA, Noc has not been shown to interact with FtsZ directly. Instead, Noc is hypothesized to block Z-ring nucleation sites by tethering the chromosome to the membrane (33). More recent, high resolution microscopy experiments suggest that rather than inhibiting Z-ring formation over the nucleoid, Noc promotes shifting of non-medial Z-ring intermediates toward midcell in a process described as “corralling”(34). Notably, neither Min nor NO are required for midcell localization of FtsZ, though each contributes to the efficiency of medial division (21).

Division in bacteria is not always medial nor occluded by the presence of nucleoid. For example, the earliest stage of *B. subtilis* sporulation is characterized by an asymmetric septation over one chromosome, resulting in two cell compartments with transiently different genetic complements (35). The larger “mother” cell contains a complete copy of the chromosome, while the smaller future spore (forespore) initially captures only a segment of the origin-proximal region of a second chromosome (36-38); bisection of the forespore-destined chromosome is avoided because the DNA is threaded through the FtsK-like ATPase, SpoIIIE, which pumps the remainder of the chromosome into the forespore (39-41). The polar division of sporulation is of considerable interest because it requires bypass of the the Min and NO systems active during vegetative growth (40, 42).

RefZ (Regulator of FtsZ) is a DNA-binding protein expressed during the early stages of sporulation (43). *refZ* is also repressed by the glucose repressor CcpA (44) and activated under conditions of phosphate limitation (45), and by the stationary phase sigma factor, SigH (46). During sporulation, RefZ binds to DNA motifs (*RBM*s) located near *oriC* (*RBM*_*O*_) and on the left and right chromosomal arms (*RBM*_*L1*_, *RBM*_*L2*_, *RBM*_*R1*_, and *RBM*_*R2*_)(47, 48). The left and right arm *RBM*s fall at the boundary demarcating the segment of chromosome localized in the forespore at the time of polar septation (47). Cells lacking *refZ* or the *RBM*s are more likely to capture DNA regions generally excluded from the forespore at the time of polar division, indicating that RefZ-*RBM* complexes somehow influence the position of the septum relative to the chromosome (47). During sporulation, a Δ*refZ* mutant also exhibits, on average, a delay in polar Z-ring formation (48) and a small shift of the septum toward midcell (49). These results suggest the Δ*refZ* mutant’s “overcapture” phenotype may be at least partially attributable to changes in chromosome organization or increased forespore dimensions that arise due to the delay.

There is likely a strong selective advantage to maintaining *RBM* positioning on the chromosome, as the arrangement of the *RBM*s on the left and right chromosome arms is remarkably conserved across the entire *Bacillus* genus (47); however, deleting *refZ* or introducing mutations into the *RBM*s that prevent RefZ binding does not reduce sporulation efficiency under standard laboratory growth conditions (47, 50). The mechanism underlying RefZ’s observed effects on FtsZ, and the selective advantage conferred to *Bacillus* by maintaining *refZ* and the *RBM*s are not known.

In this work, we performed a candidate-based screen to determine if deleting *refZ* in combination with other genes previously implicated in FtsZ regulation and/or chromosome capture would result in synthetic sporulation phenotypes. We found that combining Δ*refZ* with deletions of *ezrA, sepF, parA*, or *minD* did not detectably reduce sporulation efficiency. In contrast, reduced sporulation of a Δ*refZ* Δ*noc* mutant was evident in a plate-based assay. Using reporters of sporulation progression, we determined that the sporulation defect occurred after Spo0A activation, but prior to late sporulation. A subpopulation of Δ*refZ* Δ*noc* mutant cells failed to divide polarly or, following division, to activate the first forespore sigma factor. In addition, the Δ*refZ* Δ*noc* mutant exhibited aberrant divisions indicative of dysregulated Z-ring assembly. Our results reveal an epistatic relationship between RefZ and Noc consistent with the two proteins contributing synthetically to cell division regulation and spore development.

## RESULTS

### Deletion of *refZ* and *noc* results in a synthetic sporulation defect

RefZ promotes shifting of medial Z-rings toward the pole and the precise capture of chromosome in the forespore, yet deletion of *refZ* does not affect sporulation efficiency (47, 48), suggesting RefZ function is redundant with other factors affecting cell division and/or chromosome capture. To look for synthetic sporulation defects, we integrated a reporter of late-stage sporulation, P_*cotD*_-*lacZ*, into the chromosome at an ectopic locus. In this background, cells reaching late-stage sporulation express beta-galactosidase, resulting in accumulation of blue pigment on sporulation medium containing X-gal (Fig 1). When spotted at equivalent densities, Δ*refZ* cells turned blue at a rate indistinguishable from wildtype (Fig 1), consistent with prior results demonstrating that the Δ*refZ* mutant does not exhibit a sporulation defect (47, 48). A mutant harboring point mutations that abrogate RefZ binding at the five origin-proximal *RBM*s (47), *RBM*_*5mu*_, also sporulated indistinguishably from wildtype in the plate-based assay (Fig 1).

**Figure 1.**
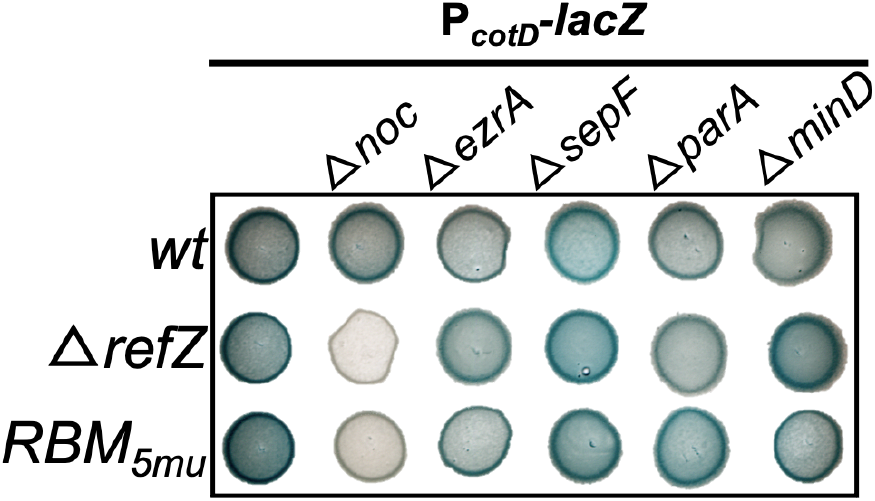
Plate-based sporulation assay based on *lacZ* expression from a late-stage sporulation promoter, P_*cotD*_. *wt* (BAM1323), Δ*noc* (BAM1321), Δ*ezrA* (BAM1563), Δ*sepF* (BAM1548), Δ*parA* (BAM1549), Δ*minD* (BAM1575), Δ*refZ* (BAM1550), Δ*refZ* Δ*noc* (BAM1546), Δ*refZ* Δ*ezrA* (BAM1559), Δ*refZ* Δ*sepF* (BAM1577), Δ*refZ* Δ*parA* (BAM1568), Δ*refZ* Δ*minD* (BAM1578), *RBM*_*5mu*_ (BAM1573), *RBM*_*5mu*_ Δ*noc* (BAM1562), *RBM*_*5mu*_ Δ*ezrA* (BAM1564), *RBM*_*5mu*_ Δ*sepF* (BAM1547), *RBM*_*5mu*_ Δ*parA* (BAM1569), *RBM*_*5mu*_ Δ*minD* (BAM1576).

Next, we assessed expression of P_*cotD*_*-lacZ* in strains harboring deletions of genes implicated in regulating FtsZ dynamics (*noc, ezrA*, and *sepF*)(7, 32, 51-53), chromosome capture (*soj*)(53-55), or both (*minD*)(52, 53). Again, each of the single deletion strains progressed in sporulation comparably to wildtype (Fig 1). Similar results were obtained when each deletion was introduced into a Δ*refZ* or *RBM*_*5mu*_ mutant background, with two exceptions: the Δ*refZ* Δ*noc* and *RBM*_*5mu*_ Δ*noc* mutants did not turn blue during the experimental time course (Fig 1), consistent with a delay or halt in sporulation progression. We conclude that RefZ and the *RBM*s contribute synthetically with Noc to support wild-type sporulation.

### Δ*refZ* Δ*noc and RBM*_*5mu*_ Δ*noc* mutants initiate sporulation

The sporulation delay observed in the Δ*refZ* Δ*noc* and *RBM*_*5mu*_ Δ*noc* double mutants could be due to failed or reduced entry into the sporulation program or to a delay or halt at any stage of sporulation prior to *cotD* expression. To investigate further, we introduced a fluorescent reporter (P_*spoIIG*_*-CFP*) activated during the earliest stage of sporulation by the sporulation master regulator, Spo0A-P (44, 56). Using P_*spoIIG*_*-CFP*, we were able to identify cells that had initiated sporulation, including cells without polar septa. Two and a half hours after resuspension in sporulation medium, cells were collected, and CFP expression was monitored by epifluorescence microscopy. The single and double mutants were indistinguishable from wildtype with respect to the number of CFP-expressing cells (Fig 2 and Fig S1), indicating that the sporulation defect in Δ*refZ* Δ*noc* and *RBM*_*5mu*_ Δ*noc* mutant cells occurs after sporulation is initiated.

**Figure 2.**
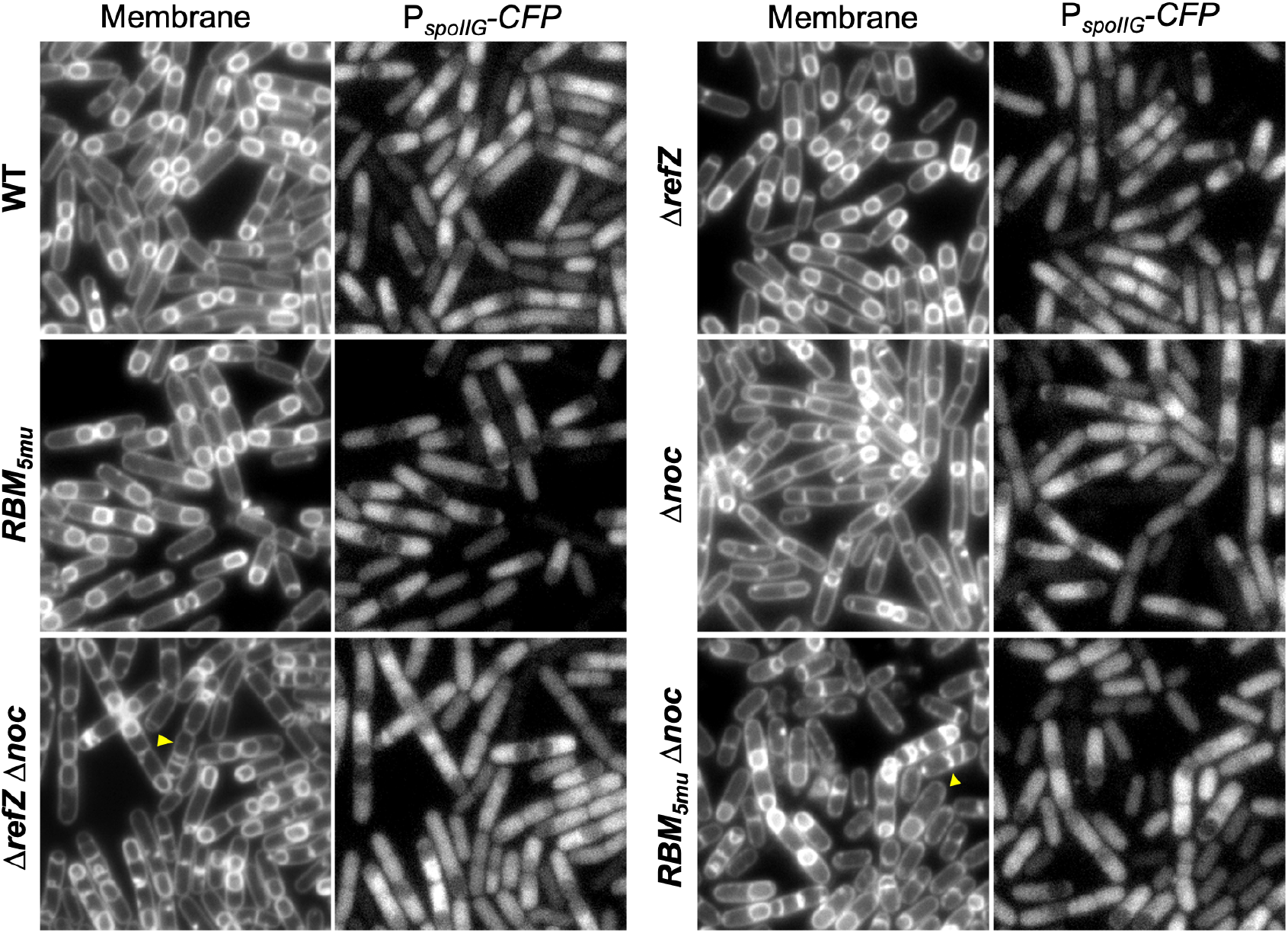
Expression of CFP from Spo0A-dependent promoter P_*spoIIG*_. Images were captured 2.5 hr following sporulation by resuspension. *wt* (BAM909), Δ*refZ* (BAM1603), *RBM*_*5mu*_ (BAM910), Δ*noc* (BAM912) Δ*refZ* Δ*noc* (BAM1604), *RBM*_*5mu*_ Δ*noc* (BAM920). CFP images scaled identically to allow for direct comparison of fluorescence.

While the double mutants displayed no qualitative delay in Spo0A-P activation, it was evident that the frequency of cells expressing CFP but lacking polar septa was increased (Fig 2 and Fig S1). To quantitate, the CFP channels of all images were scaled identically, and cells lacking polar septa were scored as expressing CFP either above or below a fixed threshold (Fig 3 and Table 1). Notably while 5-12% of wildtype and single mutant cells expressing CFP lacked septa, this value increased to 36% and 54% in the Δ*refZ* Δ*noc* and *RBM*_*5mu*_ Δ*noc* double mutants, respectively (Table 1). The increase was observed across biological and experimental replicates. We conclude that a subset of sporulating Δ*refZ* Δ*noc* and *RBM*_*5mu*_ Δ*noc* double mutants initiate sporulation, but then fail to progress to polar division during the experimental timecourse.

**Table 1.**
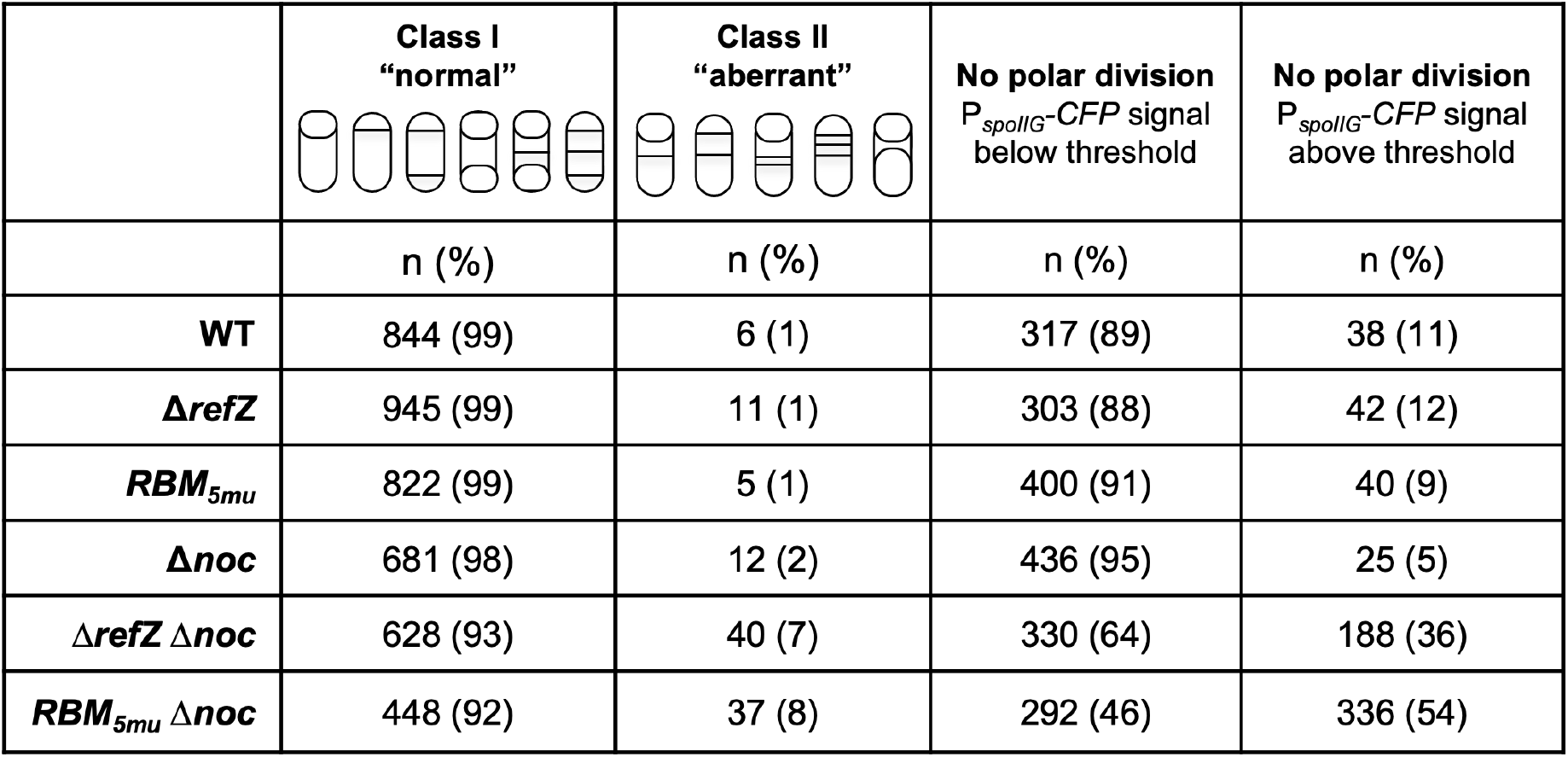
Cell division and P_*spoIIG*_-*CFP* signal classes during sporulation.

| | **Class I**<br>**"normal"** | **Class II**<br>**"aberrant"** | **No polar division**<br>$P_{spoII\bar{G}}$-CFP signal<br>below threshold | **No polar division**<br>$P_{spoII\bar{G}}$-CFP signal<br>above threshold |
| --- | --- | --- | --- | --- |
|  | n (%) | n (%) | n (%) | n (%) |
| **WT** | 844 (99) | 6 (1) | 317 (89) | 38 (11) |
| **$\Delta refZ$** | 945 (99) | 11 (1) | 303 (88) | 42 (12) |
| **$RBM_{5mu}$** | 822 (99) | 5 (1) | 400 (91) | 40 (9) |
| **$\Delta noc$** | 681 (98) | 12 (2) | 436 (95) | 25 (5) |
| **$\Delta refZ \Delta noc$** | 628 (93) | 40 (7) | 330 (64) | 188 (36) |
| **$RBM_{5mu} \Delta noc$** | 448 (92) | 37 (8) | 292 (46) | 336 (54) |

**Figure 3.**
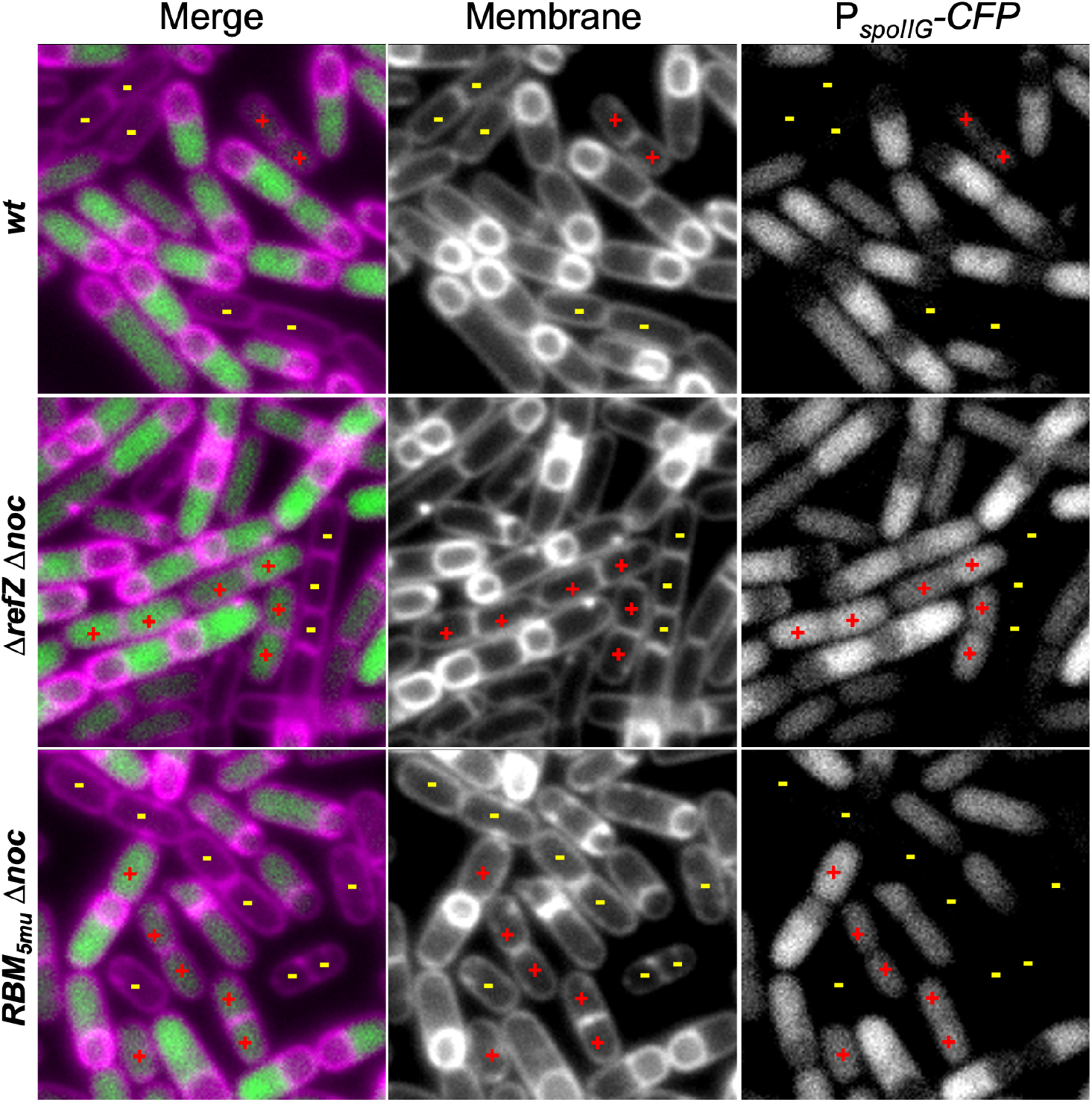
Expression of CFP from Spo0A-dependent promoter P_*spoIIG*_ in non-dividing cells. Images were captured 2.5 hr following sporulation by resuspension. *wt* (BAM909), Δ*refZ* Δ*noc* (BAM1604), *RBM*_*5mu*_ Δ*noc* (BAM920). Membranes were stained with TMA. CFP images scaled identically to allow for direct comparison of fluorescence. All images are the same magnification. Examples of non-dividing cells exhibiting CFP fluorescence below (yellow dash) and above (red plus sign) a defined threshold.

### The Δ*refZ* Δ*noc and RBM*_*5mu*_ Δ*noc* double mutants divide aberrantly during sporulation

At 2.5 hr sporulation, the majority of wildtype and single mutant cells expressing P_*spoIIG*_*-CFP* possessed an asymmetric septum or had progressed to the engulfment stage, when forespores appear rounded (Fig 2 and Fig S1). Asymmetric septa and engulfment were also observed in the Δ*refZ* Δ*noc* and *RBM*_*5mu*_ Δ*noc* mutants; however, unlike the single mutants, cells with abnormal morphological features were also readily observed (Fig 2 and Fig S1). The most frequent abnormal phenotype was a cell with two septa, one polar and one at approximately midcell of the presumed mother cell compartment (Fig 2, yellow arrowheads & Table 1). Notably, while only 1% of wild-type cells and 1-2% of the single mutants divided aberrantly (Table 1, Class II), this percentage increased to 6% and 7% in the Δ*refZ* Δ*noc* and *RBM*_*5mu*_ Δ*noc* double mutants, respectively. We conclude that there is a synthetic interaction between the activities of RefZ (likely bound at *RBM*s) and Noc that inhibits aberrant divisions during wild-type sporulation.

### The Δ*noc* Δ*refZ and RBM*_*5mu*_ Δ*noc* double mutants activate SigF

Following polar division, sporulation progression requires activation of SigF, the first forespore-specific sigma factor. To assess if the polar compartments resembling forespores activated SigF in the mutants, we monitored expression of a SigF-dependent reporter, P_*spoIIQ*_*-CFP* (57-59) using epifluorescence microscopy (Fig 4). The percentage of cells with activated SigF across the entire population of cells with forespore-like compartments was comparable for wildtype (85%), Δ*refZ* Δ*noc* (89%), and *RBM*_*5mu*_ Δ*noc* (86%)(Table 2). Consistent with our other results, only 1% of wildtype cells exhibited aberrant (Class II) divisions, while the proportion was higher for the Δ*refZ* Δ*noc* (6%) and *RBM*_*5mu*_ Δ*noc* (7%) mutants (Fig 4 and Table 2). Of cells with aberrant divisions, 100% of wildtype exhibited SigF activation compared to 39% and 56% in the Δ*refZ* Δ*noc* and *RBM*_*5mu*_ Δ*noc* mutants, respectively (Fig 4, Table 2). We conclude that Δ*refZ* Δ*noc* and *RBM*_*5mu*_ Δ*noc* cells that divide asymmetrically are still able to activate SigF in the smaller cell compartments, but with reduced frequency compared to wildtype.

**Table 2.** Cell division and P_*spoIIQ*_-*CFP* forespore signal (SigF activation) classes during sporulation.

|  | **Class I**<br>**“normal”**<br>**(Total)** | **Class II**<br>**“aberrant”**<br>**(Total)** | **Forespores**<br>**(Total)**<br>**$P_{spoIIQ}$-CFP signal**<br>**above threshold** | **Forespores**<br>**(Total)**<br>**$P_{spoIIQ}$-CFP signal**<br>**below threshold** | **Forespores**<br>**(Class II only)**<br>**$P_{spoIIQ}$-CFP signal**<br>**above threshold** | **Forespores**<br>**(Class II only)**<br>**$P_{spoIIQ}$-CFP signal**<br>**below threshold** |
| --- | --- | --- | --- | --- | --- | --- |
|  | n (%) | n (%) | n (%) | n (%) | n (%) | n (%) |
| **WT** | 655 (99) | 4 (1) | 563 (85) | 96 (15) | 4 (100) | 0 (0) |
| **$\Delta refZ$**<br>**$\Delta noc$** | 445 (93) | 36 (7) | 430 (89) | 51 (11) | 22 (61) | 14 (39) |
| **$RBM_{5mu}$**<br>**$\Delta noc$** | 528 (94) | 36 (6) | 487 (86) | 77 (14) | 16 (44) | 20 (56) |

**Figure 4.**
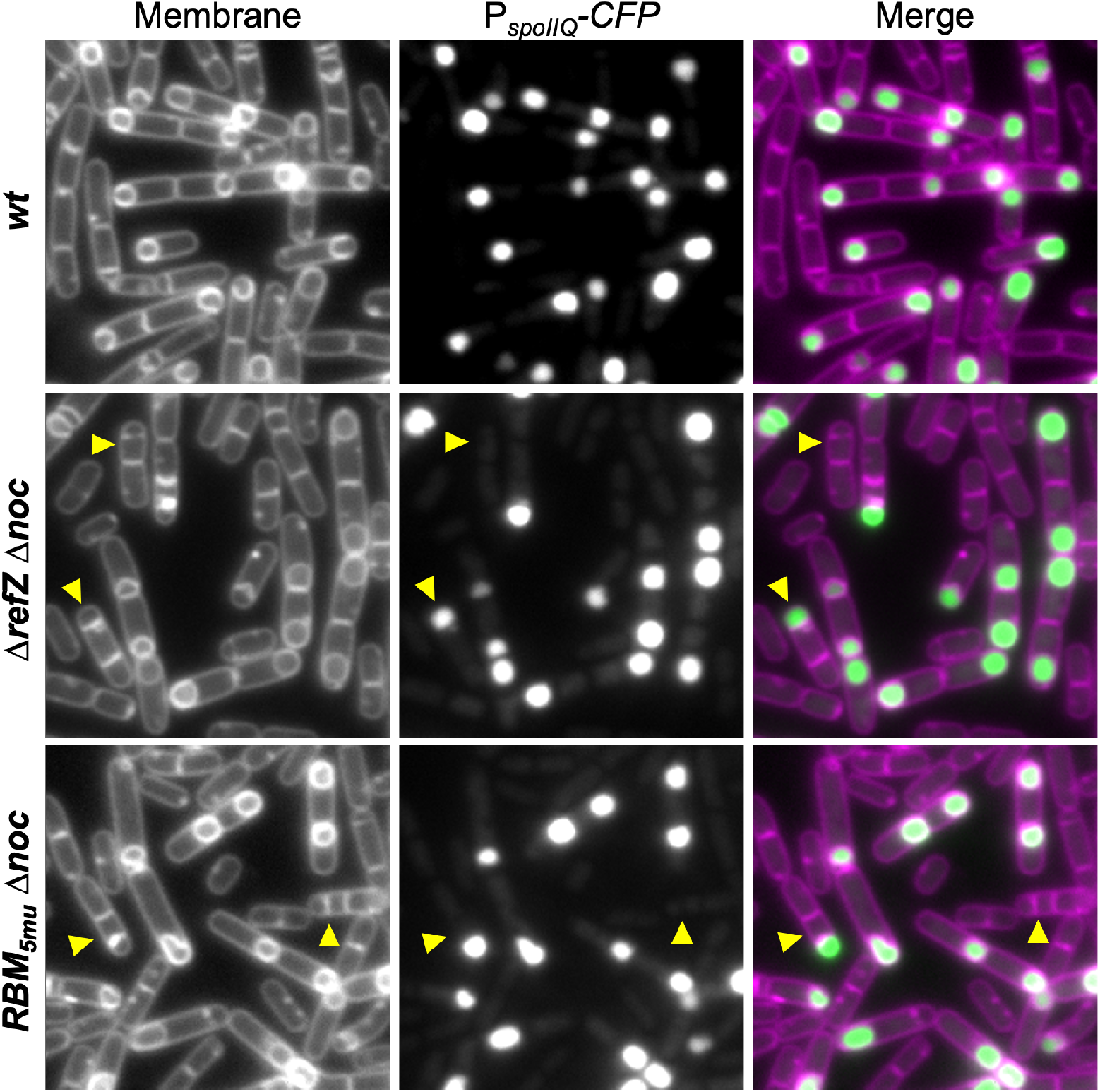
Expression from P_*spoIIQ*_*-CFP*, a SigF-dependent reporter. Images were captured 2.5 hr following sporulation by resuspension. WT (BAM1638), Δ*refZ* Δ*noc* (BAM1639), *RBM*_*5mu*_ Δ*noc* (BAM1640). Membranes were stained with TMA. CFP channels are scaled identically to allow for direct comparison. Images are identical magnification. Yellow arrowheads indicate examples of aberrantly dividing cells (Class II).

### RefZ’s division regulation activity is required for preventing aberrant septum formation in the absence of Noc

We previously identified 10 RefZ loss-of-function variants (rLOFs) that retain the ability to bind DNA but are no longer able to perturb Z-ring assembly when artificially induced during vegetative growth (60). Cells harboring the rLOF alleles in place of wild-type *refZ* miscapture DNA in the forespore indistinguishably from the Δ*refZ* mutant, suggesting that RefZ-*RBM* complexes affect chromosome capture through direct or indirect effects on FtsZ (60). We hypothesized that RefZ’s ability to modulate FtsZ activity would also be required to prevent aberrant divisions in the absence of *noc*. To test, we first replaced native *refZ* with each of the 10 rLOF alleles in a Δ*noc* background and evaluated sporulation using the plate-based P_*cotD*_*-lacZ* assay. In the presence of wild-type *noc*, each of the rLOF encoding variants supported sporulation at levels indistinguishable from wildtype (wt) or a strain encoding wild-type *refZ* linked to a chloramphenicol resistance cassette (WT - isogenic to the rLOF strains)(Fig 5). Conversely, none of the rLOF variants supported wild-type sporulation in a Δ*noc* background (Fig 5), as observed with the Δ*refZ* Δ*noc* and *RBM*_*5mu*_ Δ*noc* double mutants (Fig 1). These results suggest RefZ’s ability to affect FtsZ is also required to support wild-type sporulation in the absence of Noc.

**Figure 5.**
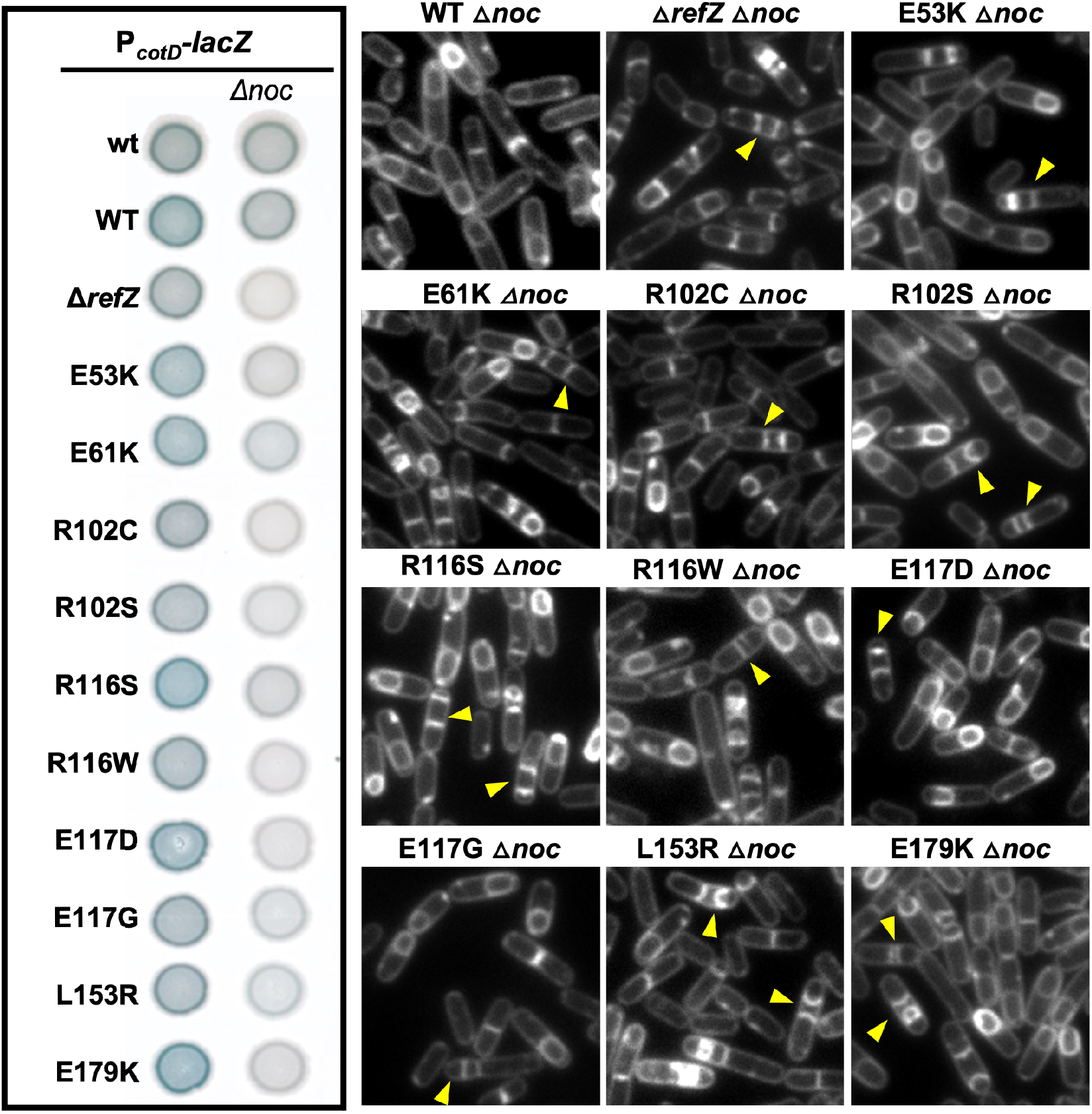
RefZ *LOFs* phenocopy Δ*refZ* with respect to supporting sporulation in the absence of *noc*. (A) Plate-based sporulation assay based on *lacZ* expression from a late-stage sporulation promoter, P_*cotD*_. Strains from left to right, top to bottom: *wt* (BAM1323), Δ*noc* (BAM1321); WT (BAM1324), WT Δ*noc* (BAM1305); Δ*refZ* (BAM1322), Δ*refZ* Δ*noc* (BAM1230); E53K (BAM1325), E53K Δ*noc* (BAM1306); E61K (BAM1326), E61K Δ*noc* (BAM1307); R102C (BAM1327), E61K Δ*noc* (BAM1308); R102S (BAM1328), R102S Δ*noc* (BAM1309); R116S (BAM1329), R116S Δ*noc* (BAM1310); R116W (BAM1330), R116W Δ*noc* (BAM1311); E117D (BAM1331), E117D Δ*noc* (BAM1312); E117G (BAM1332), E117G Δ*noc* (BAM1313); L153R (BAM1333), L153R Δ*noc* (BAM1314); E179K (BAM1334), E179K Δ*noc* (BAM1315). (B) Images were captured 3.5 hr following sporulation by resuspension. Membranes were stained with TMA. All images are at identical magnification. Yellow arrowheads indicate examples of aberrant (Class II) divisions. WT Δ*noc* (BAM1280), Δ*refZ* Δ*noc* (BAM1295), E53K Δ*noc* (BAM1281), E61K Δ*noc* (BAM1282), R102C Δ*noc* (BAM1283), R102S Δ*noc* (BAM1284), R116S Δ*noc* (BAM1285), R116W Δ*noc* (BAM1286), E117D Δ*noc* (BAM1287), E117G Δ*noc* (BAM1288), (L153R) Δ*noc* (BAM1289), E179K Δ*noc* (BAM1290).

To determine whether the sporulation defect observed in the rLOF Δ*noc* double mutants also resulted in increased aberrant divisions, we monitored division in sporulating cells using fluorescence microscopy. Aberrant divisions were rarely observed in the isogenic wild-type *refZ* strain or the rLOF mutant strains; however, when paired with Δ*noc*, each of the rLOF mutants phenocopied the Δ*refZ* Δ*noc* and *RBM*_*5mu*_ Δ*noc* double mutants (Fig 5). We conclude that the residues of RefZ that are required to affect division and support wild-type chromosome capture are also required to prevent abnormal divisions during sporulation in the absence of *noc*.

## DISCUSSION

Both RefZ and Noc are DNA-binding proteins previously implicated in FtsZ regulation. RefZ is expressed early in sporulation and facilitates both timely polar Z-ring assembly (48) and precise capture of DNA in the forespore (47, 48, 61). A Δ*refZ* mutant sporulates with wild-type efficiency, so the functional significance of RefZ activity during sporulation remains unclear (48). Noc is also expressed during sporulation but to our knowledge has not previously been associated with a sporulation function. Here we find that deleting both *noc* and *refZ* results in a synthetic sporulation defect characterized by aberrant divisions and stalled sporulation progression.

We observed several types of sporulation defects in the Δ*refZ* Δ*noc* mutant, each of which may contribute to the reduced sporulation observed in the plate-based assay. In one type, cells initiated the sporulation program, but failed to divide polarly. In a second, cells initiated sporulation and divided polarly, but failed to activate SigF, the first forespore-specific sigma factor. The largest category of aberrant cells observed divided polarly and activated SigF, but also possessed an extra septum near midcell. Some of these extra septa appear to curve at later timepoints, reminiscent of early engulfment (several examples can be seen in Fig 5). The “mother” cell chromosome may be pumped into the forespore-distal compartment in these cells. If so, the curvature might be explained if the center compartment attempted to engulf the abnormally large “twin” via residual SigE programming. We did not investigate the phenotype further in the present study, but propose the Δ*refZ* Δ*noc* mutant may be useful for interrogating models of engulfment and spore morphogenesis (62).

Genetic interactions between Noc and other proteins implicated in cell division regulation have been observed previously (63, 64). Under conditions of rapid growth, a Δ*noc* Δ*minD* mutant is filamentous and lyses, suggesting interplay between the activities of Noc and MinD. Though we lack a mechanistic understanding of how Noc and RefZ influence Z-ring formation, the fact a sporulation defect was only observed when combining Δ*refZ* with Δ*noc* (Fig 1), suggests there is some specificity to the interaction. Both RefZ and Noc spread along DNA and require DNA-binding for activity (61, 65). Transcriptomic profiling and identification of the RefZ and Noc binding sites did not reveal obvious regulons (32, 48, 51). Of note, the NO protein SlmA is also not considered to be a transcription factor in *E. coli*; however, SlmA has been shown to activate chitobiose utilization in another enteric, *Vibrio cholerae* (66). It may be informative to revisit the effects of RefZ, Noc, and SlmA on transcription under varying growth conditions using more modern methods (RNA-seq vs. microarrays) and also reexamine the regions where RefZ, Noc, and SlmA bind to look for relationships between the genes aside from location.

The *noc* gene evolved following a duplication of *spo0J* (*parB*) in the Firmicutes (67). In *B. subtilis*, Noc and Spo0J are 38% identical. Both proteins bind DNA and are regulated by CTP (65, 68-70); however, Noc, unlike Spo0J (71-74), does not appear to play a role in chromosome segregation; instead, a Δ*noc* mutant assembles aberrant Z-rings and/or divides over the chromosome, though only under conditions in which DNA replication and/or organization are perturbed (51, 75, 76). By comparison Δ*noc* mutants in *Staphylococcus aureus* sometimes assemble extra Z-rings or divide over chromosomes, but also overinitiate DNA replication (76, 77). Several lines of evidence suggest that, at least in *S. aureus*, Noc’s influence on Z-ring assembly is sensitive to nucleotide pools. First, in cells lacking *comEB* (encoding a putative CMP/dCMP deaminase), *noc* becomes essential (76). Characterized CMP/dCMP deaminases generate dCMP from dUMP, a precursor required for dTTP synthesis. Second, mutations in *dnaA* that reduce DNA replication initiation suppress Δ*noc* Δ*comEB* synthetic lethality and reduce the aberrant Z-rings associated with Δ*noc* (76). Third, Δ*noc* mutants are sensitized to DnaA overexpression compared to wildtype (76). These results suggest that Noc has activity that may buffer the cell against uncoordinated DNA replication and cell division. The reason for RefZ’s synthetic interaction with Noc remains unclear, though it is notable that the aberrant divisions and failures to progress in sporulation occur at a time in development when new rounds of DNA replication are inhibited (50, 62). Although *refZ* falls within the *spo0A* regulon, transcriptional profiling suggests that sporulation is only one context in which *refZ* is expressed (43-46, 78). Any future models for RefZ function would benefit incorporating these additional expression contexts, as well as the conservation of the *RBM*s across the genus.

## MATERIALS and METHODS

### General methods

Strains and details of strain construction can be found in the supplementary materials (Table S1 and Text S1). All *B. subtilis* strains were derived from *B. subtilis* 168. For microscopy, 25 ml cultures were grown in 250 ml baffled flasks placed in a 37°C in a shaking waterbath. *B. subtilis* transformations were carried out as previously described (79), unless otherwise indicated. Sporulation was initiated by growing cells in CH medium followed by resuspension in sporulation medium (80). *B. subtilis* selections were carried out at the following antibiotic concentrations: 100 µg/ml spectinomycin, 7.5 µg/ml chloramphenicol, 10 µg/ml kanamycin, 10 µg/ml tetracycline, and 1 µg/ml erythromycin (erm) plus 25 µg/ml lincomycin for MLS. For transformation and selection in *E. coli*, antibiotics were included at the following concentrations: 100 µg/ml ampicillin and 25 µg/ml kanamycin.

### P_cotD_-lacZ sporulation assay

For the spot plate sporulation assays, isolated colonies were used to inoculate 4 ml of DSM broth (80) and cultures were grown at 37°C in a roller drum to mid-log phase. All samples were normalized to the lowest recorded culture OD600 and 5 μl from each dilution was spotted on DSM agar plates supplemented with 40 μl/ml X-gal (5-bromo-4-chloro-3-indolyl-β-d-galactopyranoside). Plates were incubated overnight at 37°C prior to imaging with a ScanJet G4050 flatbed scanner (Hewlett Packard) using VueScan software and medium format mode. Images were processed using Adobe Photoshop (vs. 12.0) and ImageJ64 (81).

### Fluorescence microscopy

Three hundred to 500 µL samples were harvested at 6,010 × *g* for 1 min in a tabletop microcentrifuge. Supernatants were aspirated and pellets were resuspended in 3-5 µL of 1X PBS containing 0.02 mM 1-(4-(trimethylamino) phenyl)-6-phenylhexa-1,3,5-triene (TMA-DPH)(Invitrogen).Cells were mounted on glass slides with polylysine-treated coverslips. Images were captured with NIS Elements Advanced Research software (version 4.10) on a Nikon Ti-E microscope fitted with a CFI Plan Apo lambda DM 100X objective, Prior Scientific Lumen 200 Illumination system, C-FL UV-2E/C DAPI filter cube with a neutral density filter and a C-FL Cyan GFP filter cube using a CoolSNAP HQ2 monochrome camera. Images were captured for 1 s. Images were analyzed in NIS-Elements or ImageJ64 (81).

## ACKNOWLEDGEMENTS

We thank members of the Herman Lab for critical reading of the manuscript. This work was supported by a grant from the National Science Foundation to J.K.H. (MCB-1514629).

## Notes

### Competing Interest Statement

The authors have declared no competing interest.

