## Supplementary material for "RefZ and Noc act synthetically to prevent aberrant divisions during *Bacillus subtilis* sporulation": Fig S1

Short title: Septation during *B. subtilis* sporulation

Figure S1. Expression of CFP from Spo0A-dependent promoter $P_{spoIIG}$ 2.5 hr following sporulation by resuspension.
Table S1. Strains
Text S1. Strain construction

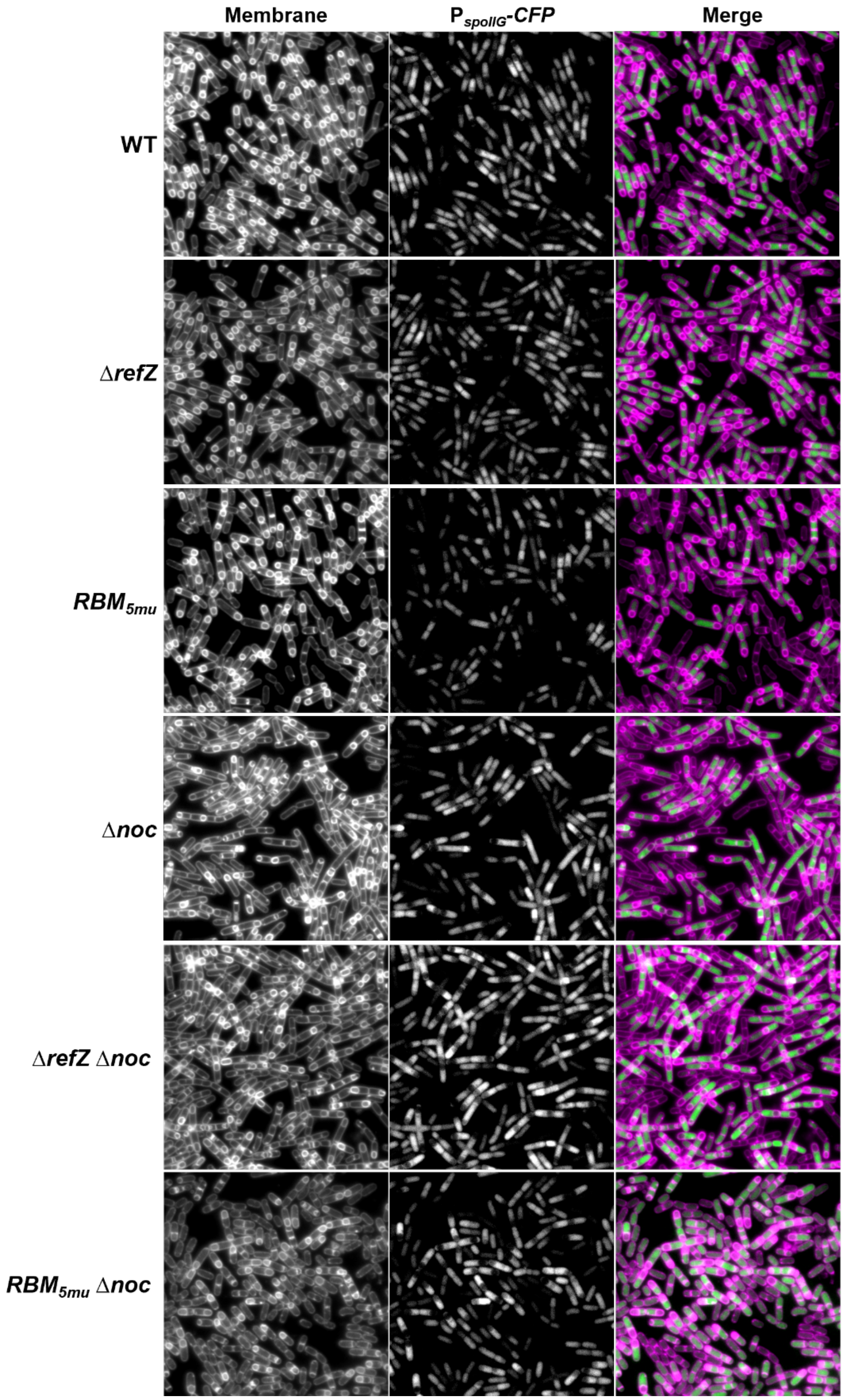

**Figure S1. Expression of CFP from Spo0A-dependent promoter $P_{spoIIG}$ 2.5 hr following sporulation by resuspension**. CFP images are scaled identically to allow for direct comparison of fluorescence. WT (BAM909), Δ*refZ* (BAM1603), $RBM_{5mu}$ (BAM910), Δ*noc* (BAM912) Δ*refZ* Δ*noc* (BAM1604), $RBM_{5mu}$ Δ*noc* (BAM920).

**Table S1. Strains**

| | | |
|---|---|---|
| ***Bacillus subtilis* 168** | Corresponds to strain stored at the *Bacillus* Genetic Stock Center under accession number 1A1. Unlike the reference genome, this strain encodes full-length SwrA. | |
| BAM043 | *minD::kan* | |
| BAM067 | *amyE::*$P_{spoIIQ}$*-cfp (cat)* | |
| BAM118 | *ezrA::kan* | |
| BAM125 | $RBM_{5mu}$*, ezrA::kan* | |
| BAM226 | $RBM_{5mu}$*, sepF::erm* | |
| BAM325 | *noc::erm* | |
| BAM469 | *Δ(soj-spo0J)::cat, pelB::spo0J (kan)* | |
| BAM908 | $RBM_{5mu}$*, noc::erm* | |
| BAM909 | *amyE::*$P_{spoIIG}$*-cfp (spec)* | Fig 2, 3, S1 |
| BAM910 | $RBM_{5mu}$*, amyE::*$P_{spoIIG}$*-cfp (spec)* | Fig 2, 3, S1 |
| BAM912 | *amyE::*$P_{spoIIG}$*-cfp (spec), noc::erm* | Fig 2, 3, S1 |
| BAM920 | $RBM_{5mu}$*, amyE::*$P_{spoIIG}$*-cfp (spec), noc::erm* | Fig 2, 3, S1 |
| BAM1265 | *refZ::refZ (WT)(cat)* | Fig 5 |
| BAM1266 | *refZ::refZ (E53K)(cat)* | Fig 5 |
| BAM1267 | *refZ::refZ (E61K)(cat)* | Fig 5 |
| BAM1268 | *refZ::refZ (R102C)(cat)* | Fig 5 |
| BAM1269 | *refZ::refZ (R102S)(cat)* | Fig 5 |
| BAM1270 | *refZ::refZ (R116S)(cat)* | Fig 5 |
| BAM1271 | *refZ::refZ (R116W)(cat)* | Fig 5 |
| BAM1272 | *refZ::refZ (E117D)(cat)* | Fig 5 |
| BAM1273 | *refZ::refZ (E117G)(cat)* | Fig 5 |
| BAM1274 | *refZ::refZ (L153R)(cat)* | Fig 5 |
| BAM1275 | *refZ::refZ (E179K)(cat)* | Fig 5 |
| BAM1280 | *refZ::refZ (WT)(cat), noc::erm* | Fig 5 |
| BAM1281 | *refZ::refZ (E53K)(cat), noc::erm* | Fig 5 |
| BAM1282 | *refZ::refZ (E61K)(cat), noc::erm* | Fig 5 |
| BAM1283 | *refZ::refZ (R102C)(cat), noc::erm* | Fig 5 |
| BAM1284 | *refZ::refZ (R102S)(cat), noc::erm* | Fig 5 |
| BAM1285 | *refZ::refZ (R116S)(cat), noc::erm* | Fig 5 |
| BAM1286 | *refZ::refZ (R116W)(cat), noc::erm* | Fig 5 |
| BAM1287 | *refZ::refZ (E117D)(cat), noc::erm* | Fig 5 |
| BAM1288 | *refZ::refZ (E117G)(cat), noc::erm* | Fig 5 |
| BAM1289 | *refZ::refZ (L153R)(cat), noc::erm* | Fig 5 |
| BAM1290 | *refZ::refZ (E179K)(cat), noc::erm* | Fig 5 |
| BAM1295 | *refZ::cat, noc::erm* | Fig 5 |
| BAM1296 | *refZ::erm* | |

| | | |
|---|---|---|
| BAM1305 | *refZ::refZ (WT)(cat), noc::erm, amyE::*$P_{cotD}$*-lacZ (spec)* | Fig 5 |
| BAM1306 | *refZ::refZ (E53K)(cat), noc::erm, amyE::*$P_{cotD}$*-lacZ (spec)* | Fig 5 |
| BAM1307 | *refZ::refZ (E61K)(cat), noc::erm, amyE::*$P_{cotD}$*-lacZ (spec)* | Fig 5 |
| BAM1308 | *refZ::refZ (R102C)(cat), noc::erm, amyE::*$P_{cotD}$*-lacZ (spec)* | Fig 5 |
| BAM1309 | *refZ::refZ (R102S)(cat), noc::erm, amyE::*$P_{cotD}$*-lacZ (spec)* | Fig 5 |
| BAM1310 | *refZ::refZ (R116S)(cat), noc::erm, amyE::*$P_{cotD}$*-lacZ (spec)* | Fig 5 |
| BAM1311 | *refZ::refZ (R116W)(cat), noc::erm, amyE::*$P_{cotD}$*-lacZ (spec)* | Fig 5 |
| BAM1312 | *refZ::refZ (E117D)(cat), noc::erm, amyE::*$P_{cotD}$*-lacZ (spec)* | Fig 5 |
| BAM1313 | *refZ::refZ (E117G)(cat), noc::erm, amyE::*$P_{cotD}$*-lacZ (spec)* | Fig 5 |
| BAM1314 | *refZ::refZ (L153R)(cat), noc::erm, amyE::*$P_{cotD}$*-lacZ (spec)* | Fig 5 |
| BAM1315 | *refZ::refZ (E179K)(cat), noc::erm, amyE::*$P_{cotD}$*-lacZ (spec)* | Fig 5 |
| BAM1320 | *refZ::cat, noc::erm, amyE::*$P_{cotD}$*-lacZ (spec)* | Fig 5 |
| BAM1321 | *noc::erm, amyE::*$P_{cotD}$*-lacZ (spec)* | Fig 1, 5 |
| BAM1322 | *refZ::cat, amyE::*$P_{cotD}$*-lacZ (spec)* | Fig 5 |
| BAM1323 | *amyE::*$P_{cotD}$*-lacZ (spec)* | Fig 1, 5 |
| BAM1324 | *refZ::refZ (WT)(cat), amyE::*$P_{cotD}$*-lacZ (spec)* | Fig 5 |
| BAM1325 | *refZ::refZ (E53K)(cat), amyE::*$P_{cotD}$*-lacZ (spec)* | Fig 5 |
| BAM1326 | *refZ::refZ (E61K)(cat), amyE::*$P_{cotD}$*-lacZ (spec)* | Fig 5 |
| BAM1327 | *refZ::refZ (R102C)(cat), amyE::*$P_{cotD}$*-lacZ (spec)* | Fig 5 |
| BAM1328 | *refZ::refZ (R102S)(cat), amyE::*$P_{cotD}$*-lacZ (spec)* | Fig 5 |
| BAM1329 | *refZ::refZ (R116S)(cat), amyE::*$P_{cotD}$*-lacZ (spec)* | Fig 5 |
| BAM1330 | *refZ::refZ (R116W)(cat), amyE::*$P_{cotD}$*-lacZ (spec)* | Fig 5 |
| BAM1331 | *refZ::refZ (E117D)(cat), amyE::*$P_{cotD}$*-lacZ (spec)* | Fig 5 |
| BAM1332 | *refZ::refZ (E117G)(cat), amyE::*$P_{cotD}$*-lacZ (spec)* | Fig 5 |
| BAM1333 | *refZ::refZ (L153R)(cat), amyE::*$P_{cotD}$*-lacZ (spec)* | Fig 5 |
| BAM1334 | *refZ::refZ (E179K)(cat), amyE::*$P_{cotD}$*-lacZ (spec)* | Fig 5 |
| BAM1339 | *ΔrefZ* | |
| BAM1359 | *ΔrefZ, noc::erm* | |
| BAM1409 | *ΔrefZ, minD::kan* | |
| BAM1529 | *ΔrefZ, ezrA::kan* | |
| BAM1546 | *ΔrefZ, noc::erm, amyE::*$P_{cotD}$*-lacZ (spec)* | Fig 1 |
| BAM1547 | $RBM_{5mu}$*, sepF::erm, amyE::*$P_{cotD}$*-lacZ (spec)* | Fig 1 |
| BAM1548 | *sepF::erm, amyE::*$P_{cotD}$*-lacZ (spec)* | Fig 1 |
| BAM1549 | *Δ(soj-spo0J)::cat, pelB::spo0J (kan), amyE::*$P_{cotD}$*-lacZ (spec)* | Fig 1 |
| BAM1550 | *ΔrefZ, amyE::*$P_{cotD}$*-lacZ (spec)* | Fig 1 |
| BAM1557 | *ΔrefZ, sepF::erm* | |
| BAM1559 | *ΔrefZ, ezrA::kan, amyE::*$P_{cotD}$*-lacZ (spec)* | Fig 1 |
| BAM1562 | $RBM_{5mu}$*, noc::erm, amyE::*$P_{cotD}$*-lacZ (spec)* | Fig 1 |
| BAM1563 | *ezrA::kan, amyE::*$P_{cotD}$*-lacZ (spec)* | Fig 1 |
| BAM1564 | $RBM_{5mu}$*, ezrA::kan, amyE::*$P_{cotD}$*-lacZ (spec)* | Fig 1 |

| | | |
|---|---|---|
| BAM1566 | *ΔrefZ, Δ(soj-spo0J)::cat, pelB::spo0J (kan)* | |
| BAM1567 | *$RBM_{5mu}$, Δ(soj-spo0J)::cat, pelB::spo0J (kan)* | |
| BAM1568 | *ΔrefZ, Δ(soj-spo0J)::cat, pelB::spo0J (kan), amyE::PcotD-lacZ (spec)* | Fig 1 |
| BAM1569 | *$RBM_{5mu}$, Δ(soj-spo0J)::cat, pelB::spo0J (kan), amyE::$P_{cotD}$-lacZ (spec)* | Fig 1 |
| BAM1573 | *$RBM_{5mu}$, amyE::$P_{cotD}$-lacZ (spec)* | Fig 1 |
| BAM1575 | *amyE::$P_{cotD}$-lacZ (spec), minD::kan* | Fig 1 |
| BAM1576 | *$RBM_{5mu}$, amyE::$P_{cotD}$-lacZ (spec), minD::kan* | Fig 1 |
| BAM1577 | *ΔrefZ, sepF::erm, amyE::$P_{cotD}$-lacZ (spec)* | Fig 1 |
| BAM1578 | *ΔrefZ, minD::kan, amyE::$P_{cotD}$-lacZ (spec)* | Fig 1 |
| BAM1600 | *refZ::tet, amyE::$P_{spoIIQ}$-cfp (cat)* | |
| BAM1601 | *$RBM_{5mu}$, amyE::$P_{spoIIQ}$-cfp (cat)* | |
| BAM1603 | *amyE::$P_{spoIIG}$-cfp (spec), refZ::tet* | Fig 2, 3 |
| BAM1604 | *amyE::$P_{spoIIG}$-cfp (spec), noc::erm, refZ::tet* | Fig 2, 3 |
| BAM1610 | *refZ::tet, amyE::$P_{spoIIQ}$-cfp (cat), noc::erm* | |
| BAM1611 | *$RBM_{5mu}$, amyE::$P_{spoIIQ}$-cfp (cat), noc::erm* | |
| BAM1638 | *amyE::$P_{spoIIQ}$-cfp (cat), rpoC-gfp (spec)* | Fig 4 |
| BAM1639 | *refZ::tet, amyE::$P_{spoIIQ}$-cfp (cat), noc::erm, rpoC-gfp (spec)* | Fig 4 |
| BAM1640 | *$RBM_{5mu}$, amyE::$P_{spoIIQ}$-cfp (cat), noc::erm, rpoC-gfp (spec)* | Fig 4 |
| BJH205 | *$RBM_{5mu}$* | (1) |
| BJH358 | *sepF::erm* (BKE15390) | (2) |

**Text S1. Strain construction**

**BAM043** [*minD::kan*] was created by transforming *Bs168* with genomic DNA from BDR2353 [*PY79 minD::kan*] selecting for growth on LB plates containing 10 µg/ml kanamycin.

**BAM067** [*amyE::PspoIIQ-cfp (cat)*] was created by transforming *Bs168* with genomic DNA from BJW342 [*PY79 refZ::tet, noc::erm, rpoC-gfp (spec), amyE::PspoIIQ-cfp (cat)*] selecting for growth on LB plates containing 7.5 µg/ml chloramphenicol.

**BAM118** [*ezrA::kan*] was created by transforming *Bs168* with genomic DNA from BJW438 [*PY79 ezrA::kan*] selecting for growth on LB plates containing 10 µg/ml kanamycin.

**BAM125** [*$RBM_{5mu}$, ezrA::kan*] was created by transforming BJH205 [*$RBM_{5mu}$*] with genomic DNA from BAM118 [*PY79 ezrA::kan*] selecting for growth on LB plates containing 10 µg/ml kanamycin.

**BAM226** [*$RBM_{5mu}$, sepF::erm*] was created by transforming BJH205 [*$RBM_{5mu}$*] with genomic DNA from BJH358 [*sepF::erm*] selecting for growth on LB plates containing 1 µg/ml erythromycin (erm) plus 25 µg/ml lincomycin (MLS).

**BAM325** [*noc::erm*] was created by transforming *Bs168* with genomic DNA from BJH144 [*PY79 noc::erm*] selecting for growth on LB plates containing 1 µg/ml erythromycin (erm) plus 25 µg/ml lincomycin (MLS).

**BAM469** [*Δ(soj-spo0J)::cat, pelB::spo0J (kan)*] was created by transforming BYD029 [*Δ(soj-spo0J)::cat*] with genomic DNA from BYD030 [*pelB::spo0J (kan)*] selecting for growth on LB plates containing 10 µg/ml kanamycin.

**BAM908** [$RBM_{5mu}$, *noc::erm*] was created by transforming BJH205 [$RBM_{5mu}$] with genomic DNA from BAM325 [*noc::erm*] selecting for growth on LB plates containing 1 µg/ml erythromycin (erm) plus 25 µg/ml lincomycin (MLS).

**BAM909** [*amyE::*$P_{spoIIG}$*-cfp (spec)*] was created by transforming *Bs168* with genomic DNA from BJH369 [*PY79 amyE::*$P_{spoIIG}$*-cfp (spec)*] selecting for growth on LB plates containing 100 µg/ml spectinomycin.

**BAM910** [$RBM_{5mu}$, *amyE::*$P_{spoIIG}$*-cfp (spec)*] was created by transforming BJH205 [$RBM_{5mu}$] with genomic DNA from BJH369 [*PY79 amyE::*$P_{spoIIG}$*-cfp (spec)*] selecting for growth on LB plates containing 100 µg/ml spectinomycin.

**BAM912** [*amyE::*$P_{spoIIG}$*-cfp (spec), noc::erm*] was created by transforming BAM909 [*amyE::*$P_{spoIIG}$*-cfp (spec)*] with genomic DNA from BAM325 [*noc::erm*] selecting for growth on LB plates containing 1 µg/ml erythromycin (erm) plus 25 µg/ml lincomycin (MLS).

**BAM920** [$RBM_{5mu}$, *amyE::*$P_{spoIIG}$*-cfp (spec), noc::erm*] was created by transforming BAM910 [$RBM_{5mu}$, *amyE::*$P_{spoIIG}$*-cfp (spec)*] with genomic DNA from BAM325 [*noc::erm*] selecting for growth on LB plates containing 1 µg/ml erythromycin (erm) plus 25 µg/ml lincomycin (MLS).

**BAM1265** [*refZ::refZ (WT)(cat)*] was created by transforming BJH247 [*refZ::tet*] with genomic DNA from BAM1006 (3) selecting for growth on LB plates containing 7.5 µg/ml chloramphenicol.

**BAM1266** [*refZ::refZ (E53K)(cat)*] was created by transforming BJH247 [*refZ::tet*] with genomic DNA from BAM1024 (3) selecting for growth on LB plates containing 7.5 µg/ml chloramphenicol.

**BAM1267** [*refZ::refZ (E61K)(cat)*] was created by transforming BJH247 [*refZ::tet*] with genomic DNA from BAM1026 (3) selecting for growth on LB plates containing 7.5 µg/ml chloramphenicol.

**BAM1268** [*refZ::refZ (R102C)(cat)*] was created by transforming BJH247 [*refZ::tet*] with genomic DNA from BAM1012 (3) selecting for growth on LB plates containing 7.5 µg/ml chloramphenicol.

**BAM1269** [*refZ::refZ (R102S)(cat)*] was created by transforming BJH247 [*refZ::tet*] with genomic DNA from BAM1014 (3) selecting for growth on LB plates containing 7.5 µg/ml chloramphenicol.

**BAM1270** [*refZ::refZ (R116S)(cat)*] was created by transforming BJH247 [*refZ::tet*] with genomic DNA from BAM1020 (3) selecting for growth on LB plates containing 7.5 µg/ml chloramphenicol.

**BAM1271** [*refZ::refZ (R116W)(cat)*] was created by transforming BJH247 [*refZ::tet*] with genomic DNA from BAM1018 (3) selecting for growth on LB plates containing 7.5 µg/ml chloramphenicol.

**BAM1272** [*refZ::refZ (E117D)(cat)*] was created by transforming BJH247 [*refZ::tet*] with genomic DNA from BAM1022 ((3) selecting for growth on LB plates containing 7.5 µg/ml chloramphenicol.

**BAM1273** [*refZ::refZ (E117G)(cat)*] was created by transforming BJH247 [*refZ::tet*] with genomic DNA from BAM1010 (3) selecting for growth on LB plates containing 7.5 µg/ml chloramphenicol.

**BAM1274** [*refZ::refZ (L153R)(cat)*] was created by transforming BJH247 [*refZ::tet*] with genomic DNA from BAM1016 (3) for growth on LB plates containing 7.5 µg/ml chloramphenicol.

**BAM1275** [*refZ::refZ (E179K)(cat)*] was created by transforming BJH247 [*refZ::tet*] with genomic DNA from BAM1008 (3) selecting for growth on LB plates containing 7.5 µg/ml chloramphenicol.

**BAM1280** [*refZ::refZ (WT)(cat), noc::erm*] was created by transforming BAM1265 [*refZ::refZ (WT)(cat)*] with genomic DNA from BAM325 [*noc::erm*] selecting for growth on LB plates containing 1 µg 10 µg/ml erythromycin (erm) plus 25 µg 10 µg/ml lincomycin (MLS).

**BAM1281** [*refZ::refZ (E53K)(cat), noc::erm*] was created by transforming BAM1266 [*refZ::refZ (E53K)(cat)*] with genomic DNA from BAM325 [*noc::erm*] selecting for growth on LB plates containing 1 µg/ml erythromycin (erm) plus 25 µg/ml lincomycin (MLS).

**BAM1282** [*refZ::refZ (E61K)(cat), noc::erm*] was created by transforming BAM1267 [*refZ::refZ (E61K)(cat)*] with genomic DNA from BAM325 [*noc::erm*] selecting for growth on LB plates containing 1 µg/ml erythromycin (erm) plus 25 µg/ml lincomycin (MLS).

**BAM1283** [*refZ::refZ (R102C)(cat), noc::erm*] was created by transforming BAM1268 [*refZ::refZ (R102C)(cat)*] with genomic DNA from BAM325 [*noc::erm*] selecting for growth on LB plates containing 1 µg/ml erythromycin (erm) plus 25 µg/ml lincomycin (MLS).

**BAM1284** [*refZ::refZ (R102S)(cat), noc::erm*] was created by transforming BAM1269 [*refZ::refZ (R102S)(cat)*] with genomic DNA from BAM325 [*noc::erm*] selecting for growth on LB plates containing 1 µg/ml erythromycin (erm) plus 25 µg/ml lincomycin (MLS).

**BAM1285** [*refZ::refZ (R116S)(cat), noc::erm*] was created by transforming BAM1270 [*refZ::refZ (R116S)(cat)*] with genomic DNA from BAM325 [*noc::erm*] selecting for growth on LB plates containing 1 µg/ml erythromycin (erm) plus 25 µg/ml lincomycin (MLS).

**BAM1286** [*refZ::refZ (R116W)(cat), noc::erm*] was created by transforming BAM1271 [*refZ::refZ (R116W)(cat)*] with genomic DNA from BAM325 [*noc::erm*] selecting for growth on LB plates containing 1 µg/ml erythromycin (erm) plus 25 µg/ml lincomycin (MLS).

**BAM1287** [*refZ::refZ (E117D)(cat), noc::erm*] was created by transforming BAM1272 [*refZ::refZ (E117D)(cat)*] with genomic DNA from BAM325 [*noc::erm*] selecting for growth on LB plates containing 1 µg/ml erythromycin (erm) plus 25 µg/ml lincomycin (MLS).

**BAM1288** [*refZ::refZ (E117G)(cat), noc::erm*] was created by transforming BAM1273 [*refZ::refZ (E117G)(cat)*] with genomic DNA from BAM325 [*noc::erm*] selecting for growth on LB plates containing 1 µg/ml erythromycin (erm) plus 25 µg/ml lincomycin (MLS).

**BAM1289** [*refZ::refZ (L153R)(cat), noc::erm*] was created by transforming BAM1274 [*refZ::refZ (L153R)(cat)*] with genomic DNA from BAM325 [*noc::erm*] selecting for growth on LB plates containing 1 µg/ml erythromycin (erm) plus 25 µg/ml lincomycin (MLS).

**BAM1290** [*refZ::refZ (E179K)(cat), noc::erm*] was created by transforming BAM1275 [*refZ::refZ (E179K)(cat)*] with genomic DNA from BAM325 [*noc::erm*] selecting for growth on LB plates containing 1 µg/ml erythromycin (erm) plus 25 µg/ml lincomycin (MLS).

**BAM1295** [*refZ::cat, noc::erm*] was created by transforming BJH255 [*refZ::cat*] with genomic DNA from BAM325 [*noc::erm*] selecting for growth on LB plates containing 1 µg/ml erythromycin (erm) plus 25 µg/ml lincomycin (MLS).

**BAM1296** [*refZ::erm*] was created by transforming *Bs168* with genomic DNA from BKE29630 [*refZ::erm*] (2) selecting for growth on LB plates containing 1 µg/ml erythromycin (erm) plus 25 µg/ml lincomycin (MLS).

**BAM1305** [*refZ::refZ (WT)(cat), noc::erm, amyE::*$P_{cotD}$*-lacZ (spec)*] was created by transforming BAM1280 [*refZ::refZ (WT)(cat)*] with genomic DNA from BJH323 [*amyE::*$P_{cotD}$*-lacZ (spec)*] selecting for growth on LB plates containing 100 µg/ml spectinomycin.

**BAM1306** [*refZ::refZ (E53K)(cat), noc::erm, amyE::*$P_{cotD}$*-lacZ (spec)*] was created by transforming BAM1281 [*refZ::refZ (E53K)(cat)*] with genomic DNA from BJH323 [*amyE::*$P_{cotD}$*-lacZ (spec)*] selecting for growth on LB plates containing 100 µg/ml spectinomycin.

**BAM1307** [*refZ::refZ (E61K)(cat), noc::erm, amyE::*$P_{cotD}$*-lacZ (spec)*] was created by transforming BAM1282 [*refZ::refZ (E61K)(cat)*] with genomic DNA from BJH323 [*amyE::*$P_{cotD}$*-lacZ (spec)*] selecting for growth on LB plates containing 100 µg/ml spectinomycin.

**BAM1308** [*refZ::refZ (R102C)(cat), noc::erm, amyE::*$P_{cotD}$*-lacZ (spec)*] was created by transforming BAM1283 [*refZ::refZ (R012C)(cat)*] with genomic DNA from BJH323 [*amyE::*$P_{cotD}$*-lacZ (spec)*] selecting for growth on LB plates containing 100 µg/ml spectinomycin.

**BAM1309** [*refZ::refZ (R102S)(cat), noc::erm, amyE::*$P_{cotD}$*-lacZ (spec)*] was created by transforming BAM1284 [*refZ::refZ (R102S)(cat)*] with genomic DNA from BJH323 [*amyE::*$P_{cotD}$*-lacZ (spec)*] selecting for growth on LB plates containing 100 µg/ml spectinomycin.

**BAM1310** [*refZ::refZ (R116S)(cat), noc::erm, amyE::*$P_{cotD}$*-lacZ (spec)*] was created by transforming BAM1285 [*refZ::refZ (R116S)(cat)*] with genomic DNA from BJH323 [*amyE::*$P_{cotD}$*-lacZ (spec)*] selecting for growth on LB plates containing 100 µg/ml spectinomycin.

**BAM1311** [*refZ::refZ (R116W)(cat), noc::erm, amyE::*$P_{cotD}$*-lacZ (spec)*] was created by transforming BAM1286 [*refZ::refZ (R116W)(cat)*] with genomic DNA from BJH323 [*amyE::*$P_{cotD}$*-lacZ (spec)*] selecting for growth on LB plates containing 100 µg/ml spectinomycin.

**BAM1312** [*refZ::refZ (E117D)(cat), noc::erm, amyE::*$P_{cotD}$*-lacZ (spec)*] was created by transforming BAM1287 [*refZ::refZ (E117D)(cat)*] with genomic DNA from BJH323 [*amyE::*$P_{cotD}$*-lacZ (spec)*] selecting for growth on LB plates containing 100 µg/ml spectinomycin.

**BAM1313** [*refZ::refZ (E117G)(cat), noc::erm, amyE::*$P_{cotD}$*-lacZ (spec)*] was created by transforming BAM1288 [*refZ::refZ (E117G)(cat)*] with genomic DNA from BJH323 [*amyE::*$P_{cotD}$*-lacZ (spec)*] selecting for growth on LB plates containing 100 µg/ml spectinomycin.

**BAM1314** [*refZ::refZ (L153R)(cat), noc::erm, amyE::*$P_{cotD}$*-lacZ (spec)*] was created by transforming BAM1289 [*refZ::refZ (L153R)(cat)*] with genomic DNA from BJH323 [*amyE::*$P_{cotD}$*-lacZ (spec)*] selecting for growth on LB plates containing 100 µg/ml spectinomycin.

**BAM1315** [*refZ::refZ (E179K)(cat), noc::erm, amyE::*$P_{cotD}$*-lacZ (spec)*] was created by transforming BAM1290 [*refZ::refZ (E179K)(cat)*] with genomic DNA from BJH323 [*amyE::*$P_{cotD}$*-lacZ (spec)*] selecting for growth on LB plates containing 100 µg/ml spectinomycin.

**BAM1320** [*refZ::cat, noc::erm, amyE::*$P_{cotD}$*-lacZ (spec)*] was created by transforming BAM1295 [*refZ::cat, noc::erm*] with genomic DNA from BJH323 [*amyE::*$P_{cotD}$*-lacZ (spec)*] selecting for growth on LB plates containing 100 µg/ml spectinomycin.

**BAM1321** [*noc::erm, amyE::*$P_{cotD}$*-lacZ (spec)*] was created by transforming BAM325 [*noc::erm*] with genomic DNA from BJH323 [*amyE::*$P_{cotD}$*-lacZ (spec)*] selecting for growth on LB plates containing 100 µg/ml spectinomycin.

**BAM1322** [*refZ::cat*, *amyE::*$P_{cotD}$*-lacZ (spec)*] was created by transforming BJH255 [*refZ::cat*] with genomic DNA from BJH323 [*amyE::*$P_{cotD}$*-lacZ (spec)*] selecting for growth on LB plates containing 100 µg/ml spectinomycin.

**BAM1323** [*amyE::*$P_{cotD}$*-lacZ (spec)*] was created by transforming *Bs168* with genomic DNA from BJH323 [*amyE::*$P_{cotD}$*-lacZ (spec)*] selecting for growth on LB plates containing 100 µg/ml spectinomycin.

**BAM1324** [*refZ::refZ (WT)(cat)*, *amyE::*$P_{cotD}$*-lacZ (spec)*] was created by transforming BAM1265 [*refZ::refZ (WT)(cat)*] with genomic DNA from BJH323 [*amyE::*$P_{cotD}$*-lacZ (spec)*] selecting for growth on LB plates containing 100 µg/ml spectinomycin.

**BAM1325** [*refZ::refZ (E53K)(cat)*, *amyE::*$P_{cotD}$*-lacZ (spec)*] was created by transforming BAM1266 [*refZ::refZ (WT)(cat)*] with genomic DNA from BJH323 [*amyE::*$P_{cotD}$*-lacZ (spec)*] selecting for growth on LB plates containing 100 µg/ml spectinomycin.

**BAM1326** [*refZ::refZ (E61K)(cat)*, *amyE::*$P_{cotD}$*-lacZ (spec)*] was created by transforming BAM1267 [*refZ::refZ (WT)(cat)*] with genomic DNA from BJH323 [*amyE::*$P_{cotD}$*-lacZ (spec)*] selecting for growth on LB plates containing 100 µg/ml spectinomycin.

**BAM1327** [*refZ::refZ (R102C)(cat)*, *amyE::*$P_{cotD}$*-lacZ (spec)*] was created by transforming BAM1268 [*refZ::refZ (WT)(cat)*] with genomic DNA from BJH323 [*amyE::*$P_{cotD}$*-lacZ (spec)*] selecting for growth on LB plates containing 100 µg/ml spectinomycin.

**BAM1328** [*refZ::refZ (R102S)(cat)*, *amyE::*$P_{cotD}$*-lacZ (spec)*] was created by transforming BAM1269 [*refZ::refZ (WT)(cat)*] with genomic DNA from BJH323 [*amyE::*$P_{cotD}$*-lacZ (spec)*] selecting for growth on LB plates containing 100 µg/ml spectinomycin.

**BAM1329** [*refZ::refZ (R116S)(cat)*, *amyE::*$P_{cotD}$*-lacZ (spec)*] was created by transforming BAM1270 [*refZ::refZ (WT)(cat)*] with genomic DNA from BJH323 [*amyE::*$P_{cotD}$*-lacZ (spec)*] selecting for growth on LB plates containing 100 µg/ml spectinomycin.

**BAM1330** [*refZ::refZ (R116W)(cat)*, *amyE::*$P_{cotD}$*-lacZ (spec)*] was created by transforming BAM1271 [*refZ::refZ (WT)(cat)*] with genomic DNA from BJH323 [*amyE::*$P_{cotD}$*-lacZ (spec)*] selecting for growth on LB plates containing 100 µg/ml spectinomycin.

**BAM1331** [*refZ::refZ (E117D)(cat)*, *amyE::*$P_{cotD}$*-lacZ (spec)*] was created by transforming BAM1272 [*refZ::refZ (WT)(cat)*] with genomic DNA from BJH323 [*amyE::*$P_{cotD}$*-lacZ (spec)*] selecting for growth on LB plates containing 100 µg/ml spectinomycin.

**BAM1332** [*refZ::refZ (E117G)(cat)*, *amyE::*$P_{cotD}$*-lacZ (spec)*] was created by transforming BAM1273 [*refZ::refZ (WT)(cat)*] with genomic DNA from BJH323 [*amyE::*$P_{cotD}$*-lacZ (spec)*] selecting for growth on LB plates containing 100 µg/ml spectinomycin.

**BAM1333** [*refZ::refZ (L153R)(cat)*, *amyE::*$P_{cotD}$*-lacZ (spec)*] was created by transforming BAM1274 [*refZ::refZ (WT)(cat)*] with genomic DNA from BJH323 [*amyE::*$P_{cotD}$*-lacZ (spec)*] selecting for growth on LB plates containing 100 µg/ml spectinomycin.

**BAM1334** [*refZ::refZ (E179K)(cat)*, *amyE::*$P_{cotD}$*-lacZ (spec)*] was created by transforming BAM1275 [*refZ::refZ (WT)(cat)*] with genomic DNA from BJH323 [*amyE::*$P_{cotD}$*-lacZ (spec)*] selecting for growth on LB plates containing 100 µg/ml spectinomycin.

BAM1339 [*ΔrefZ*] was created by transforming BAM1296 [*refZ::erm*] with pDR244 (Cre recombinase plasmid)(2) selecting for growth on LB plates containing 100 µg/ml spectinomycin at 30ºC (permissive). Isolated colonies were propagated on LB plates containing 100 µg/ml spectinomycin at 30ºC, on LB plates at the non-permissive temperature (42ºC), and on LB plates containing MLS (37ºC). Clones were streaked for isolation on LB, spec100, and MLS and a spec and erm sensitive clone was stored.

**BAM1359** [*ΔrefZ, noc::erm*] was created by transforming BAM1339 [*ΔrefZ*] with genomic DNA from BAM325 [*noc::erm*] selecting for growth on LB plates containing 1 µg/ml erythromycin (erm) plus 25 µg/ml lincomycin (MLS).

**BAM1409** [*ΔrefZ, minD::kan*] was created by transforming BAM1339 [*ΔrefZ*] with genomic DNA from BAM043 [*minD::kan*] selecting for growth on LB plates containing 10 µg/ml kanamycin.

**BAM1529** [*ΔrefZ, ezrA::kan*] was created by transforming BAM1339 [*ΔrefZ*] with genomic DNA from BAM118 [*PY79 ezrA::kan*] selecting for growth on LB plates containing 10 µg/ml kanamycin.

**BAM1546** [*ΔrefZ, noc::erm, amyE::*$P_{cotD}$*-lacZ (spec)*] was created by transforming BAM1359 [*ΔrefZ, noc::erm*] with genomic DNA from BJH323 [*amyE::*$P_{cotD}$*-lacZ (spec)*] selecting for growth on LB plates containing 100 µg/ml spectinomycin.

**BAM1547** [*RBM*$_{5mu}$*, sepF::erm, amyE::*$P_{cotD}$*-lacZ (spec)*] was created by transforming BAM226 [*RBM*$_{5mu}$*, sepF::erm*] with genomic DNA from BJH323 [*amyE::*$P_{cotD}$*-lacZ (spec)*] selecting for growth on LB plates containing 100 µg/ml spectinomycin.

**BAM1548** [*sepF::erm, amyE::*$P_{cotD}$*-lacZ (spec)*] was created by transforming BJH358 [*sepF::erm*] with genomic DNA from BJH323 [*amyE::*$P_{cotD}$*-lacZ (spec)*] selecting for growth on LB plates containing 100 µg/ml spectinomycin.

**BAM1549** [*Δ(soj-spo0J)::cat, pelB::spo0J (kan), amyE::*$P_{cotD}$*-lacZ (spec)*] was created by transforming BAM469 [*Δ(soj-spo0J)::cat, pelB::spo0J (kan)*] with genomic DNA from BJH323 [*amyE::*$P_{cotD}$*-lacZ (spec)*] selecting for growth on LB plates containing 100 µg/ml spectinomycin.

**BAM1550** [*ΔrefZ, amyE::*$P_{cotD}$*-lacZ (spec)*] was created by transforming BAM1339 [*ΔrefZ*] with genomic DNA from BJH323 [*amyE::*$P_{cotD}$*-lacZ (spec)*] selecting for growth on LB plates containing 100 µg/ml spectinomycin.

**BAM1557** [*ΔrefZ, sepF::erm*] was created by transforming BAM1339 [*ΔrefZ*] with genomic DNA from BJH358 [*sepF::erm*] selecting for growth on LB plates containing

**BAM1559** [*ΔrefZ, ezrA::kan, amyE::*$P_{cotD}$*-lacZ (spec)*] was created by transforming BAM1529 [*ΔrefZ, ezrA::kan*] with genomic DNA from BJH323 [*amyE::*$P_{cotD}$*-lacZ (spec)*] selecting for growth on LB plates containing 100 µg/ml spectinomycin.

**BAM1562** [*RBM*$_{5mu}$*, noc::erm, amyE::*$P_{cotD}$*-lacZ (spec)*] was created by transforming BAM908 [*RBM*$_{5mu}$*, noc::erm*] with genomic DNA from BJH323 [*amyE::*$P_{cotD}$*-lacZ (spec)*] selecting for growth on LB plates containing 100 µg/ml spectinomycin.

**BAM1563** [*ezrA::kan, amyE::*$P_{cotD}$*-lacZ (spec)*] was created by transforming BAM118 [*ezrA::kan*] with genomic DNA from BJH323 [*amyE::*$P_{cotD}$*-lacZ (spec)*] selecting for growth on LB plates containing 100 µg/ml spectinomycin.

**BAM1564** [*RBM*$_{5mu}$*, ezrA::kan, amyE::*$P_{cotD}$*-lacZ (spec)*] was created by transforming BAM125 [*RBM*$_{5mu}$*, ezrA::kan*] with genomic DNA from BJH323 [*amyE::*$P_{cotD}$*-lacZ (spec)*] selecting for growth on LB plates containing 100 µg/ml spectinomycin.
**BAM1568** [*ΔrefZ, Δ(soj-spo0J)::cat, pelB::spo0J (kan), amyE::PcotD-lacZ (spec)*] was created by transforming BAM1566 [*ΔrefZ, Δ(soj-spo0J)::cat, pelB::spo0J (kan)*] with genomic DNA from BJH323 [*amyE::*$P_{cotD}$*-lacZ (spec)*] selecting for growth on LB plates containing 100 µg/ml spectinomycin.

**BAM1569** [*RBM*$_{5mu}$*, Δ(soj-spo0J)::cat, pelB::spo0J (kan), amyE::*$P_{cotD}$*-lacZ (spec)*] was created by transforming BAM1567 [*RBM*$_{5mu}$*, Δ(soj-spo0J)::cat, pelB::spo0J (kan)*] with genomic DNA from BJH323 [*amyE::*$P_{cotD}$*-lacZ (spec)*] selecting for growth on LB plates containing 100 µg/ml spectinomycin.

**BAM1573** [*$RBM_{5mu}$, amyE::$P_{cotD}$-lacZ (spec)*] was created by transforming BJH205 [*$RBM_{5mu}$*] with genomic DNA from BJH323 [*amyE::$P_{cotD}$-lacZ (spec)*] selecting for growth on LB plates containing 100 µg/ml spectinomycin.

**BAM1575** [*amyE::$P_{cotD}$-lacZ (spec), minD::kan*] was created by transforming BAM1323 [*amyE::$P_{cotD}$-lacZ (spec)*] with genomic DNA from BAM043 [*minD::kan*] selecting for growth on LB plates containing 10 µg/ml kanamycin.

**BAM1576** [*$RBM_{5mu}$, amyE::$P_{cotD}$-lacZ (spec), minD::kan*] was created by transforming BAM1573 [*$RBM_{5mu}$, amyE::$P_{cotD}$-lacZ (spec)*] with genomic DNA from BAM043 [*minD::kan*] selecting for growth on LB plates containing 10 µg/ml kanamycin.

**BAM1577** [*ΔrefZ, sepF::erm, amyE::$P_{cotD}$-lacZ (spec)*] was created by transforming BAM1557 [*ΔrefZ, sepF::erm*] with genomic DNA from BJH323 [*amyE::$P_{cotD}$-lacZ (spec)*] selecting for growth on LB plates containing 100 µg/ml spectinomycin.

**BAM1578** [*ΔrefZ, minD::kan, amyE::$P_{cotD}$-lacZ (spec)*] was created by transforming BAM1409 [*ΔrefZ, minD::kan*] with genomic DNA from BJH323 [*amyE::$P_{cotD}$-lacZ (spec)*] selecting for growth on LB plates containing 100 µg/ml spectinomycin.

**BAM1600** [*refZ::tet, amyE::$P_{spoIIQ}$-cfp (cat)*] was created by transforming BJH247 [*refZ::tet*] with genomic DNA from BAM067 [*amyE::$P_{spoIIQ}$-cfp (cat)*] selecting for growth on LB plates containing 7.5 µg/ml chloramphenicol.

**BAM1601** [*$RBM_{5mu}$, amyE::$_{PspoIIQ}$-cfp (cat)*] was created by transforming BJH205 [*$RBM_{5mu}$*] with genomic DNA from BAM067 [*amyE::$P_{spoIIQ}$-cfp (cat)*] selecting for growth on LB plates containing 7.5 µg/ml chloramphenicol.

**BAM1603** [*amyE::$P_{spoIIG}$-cfp (spec), refZ::tet*] was created by transforming BAM909 [*amyE::$P_{spoIIG}$-cfp (spec)*] with genomic DNA from BJH247 [*refZ::tet*] selecting for growth on LB plates containing 10 µg/ml tetracycline.

**BAM1604** [*amyE::$P_{spoIIG}$-cfp (spec), noc::erm, refZ::tet*] was created by transforming BAM912 [*amyE::$P_{spoIIG}$-cfp (spec), noc::erm*] with genomic DNA from BJH247 [*refZ::tet*] selecting for growth on LB plates containing 10 µg/ml tetracycline.

**BAM1610** [*refZ::tet, amyE::$P_{spoIIQ}$-cfp (cat), noc::erm*] was created by transforming BAM1600 [*refZ::tet, amyE::$P_{spoIIQ}$-cfp (cat)*] with genomic DNA from BAM325 [*noc::erm*] selecting for growth at 30ºC on LB plates containing 1 µg/ml erythromycin (erm) plus 25 µg/ml lincomycin (MLS).

**BAM1611** [*$RBM_{5mu}$, amyE::$_{PspoIIQ}$-cfp (cat), noc::erm*] was created by transforming BAM1601 [*$RBM_{5mu}$, amyE::$_{PspoIIQ}$-cfp (cat)*] with genomic DNA from BAM325 [*noc::erm*] selecting for growth at 30ºC on LB plates containing 1 µg/ml erythromycin (erm) plus 25 µg/ml lincomycin (MLS).

**BAM1638** [*amyE::*$P_{spoIIQ}$*-cfp (cat), rpoC-gfp (spec)*] was created by transforming BAM067 [*amyE::*$P_{spoIIQ}$*-cfp (cat)*] with genomic DNA from BJW342 [*PY79 refZ::tet, noc::erm, rpoC-gfp (spec), amyE::PspoIIQ-cfp (cat)*] selecting for growth at 30ºC on LB plates containing 100 µg/ml spectinomycin.

**BAM1639** [*refZ::tet, amyE::*$P_{spoIIQ}$*-cfp (cat), noc::erm, rpoC-gfp (spec)*] was created by transforming BAM1610 [*refZ::tet, amyE::*$P_{spoIIG}$*-cfp (spec), noc::erm*] with genomic DNA from BJW342 [*PY79 refZ::tet, noc::erm, rpoC-gfp (spec), amyE::*$P_{spoIIQ}$*-cfp (cat)*] selecting for growth at 30ºC on LB plates containing 100 µg/ml spectinomycin.

**BAM1640** [$RBM_{5mu}$*, amyE::*$_{PspoIIQ}$*-cfp (cat), noc::erm, rpoC-gfp (spec)*] was created by transforming BAM1611 [$RBM_{5mu}$*, amyE::*$P_{spoIIG}$*-cfp (spec), noc::erm*] with genomic DNA from BJW342 [*PY79 refZ::tet, noc::erm, rpoC-gfp (spec), amyE::*$P_{spoIIQ}$*-cfp (cat)*] selecting for growth at 30ºC on LB plates containing 100 µg/ml spectinomycin.
